# Phosphate is the third nutrient monitored by TOR in *Candida albicans* and provides a target for fungal-specific indirect TOR inhibition

**DOI:** 10.1101/142745

**Authors:** Ning-Ning Liu, Peter Flanagan, Jumei Zeng, Niketa Jani, Maria E. Cardenas, Gary P. Moran, Julia R. Köhler

## Abstract

The TOR pathway regulates morphogenesis and responses to host cells in the fungal pathogen *Candida albicans*. Eukaryotic TOR complex 1 (TORC1) induces growth and proliferation in response to nitrogen and carbon source availability. Our unbiased genetic approach seeking new components of TORC1 signaling in *C. albicans* revealed that the phosphate transporter Pho84 is required for normal TORC1 activity. We found that mutants in *PHO84* are hypersensitive to rapamycin and, in response to phosphate feeding, generate less phosphorylated ribosomal protein S6 (P-S6) than wild type. The small GTPase Gtr1, a component of the TORC1-activating EGO complex, links Pho84 to TORC1. Mutants in Gtr1, but not in another TORC1-activating GTPase, Rhb1, are defective in the P-S6 response to phosphate. Overexpression of Gtr1 and of a constitutively active Gtr1^Q67L^ mutant suppress TORC1-related defects. In *S. cerevisiae pho84* mutants, constitutively active Gtr1 suppresses a TORC1 signaling defect but does not rescue rapamycin hypersensitivity. Hence connections from phosphate homeostasis to TORC1 may differ between *C. albicans* and *S. cerevisiae*. The converse direction of signaling, from TORC1 to the phosphate homeostasis (PHO) regulon, previously observed in *S. cerevisiae*, was genetically demonstrated in *C. albicans* using conditional *TOR1* alleles. A small molecule inhibitor of Pho84, an FDA-approved drug, inhibits TORC1 signaling and potentiates the activity of the antifungals amphotericin B and micafungin. Anabolic TORC1-dependent processes require significant amounts of phosphate. Our study demonstrates that phosphate availability is monitored and also controlled by TORC1, and that TORC1 can be indirectly targeted by inhibiting Pho84.

**Significance:** The human fungal pathogen *Candida albicans* uses the TOR signaling pathway to contend with varying host environments and thereby regulate cell growth. Seeking novel components of the *C. albicans* TOR pathway we identified a cell-surface phosphate importer, Pho84, and its molecular link to TOR complex 1 (TORC1). Since phosphorus is a critical element for anabolic processes like DNA replication, ribosome biogenesis, translation and membrane biosynthesis, TORC1 monitors its availability in regulating these processes. By depleting the central kinase in the TORC1 pathway, we showed that TORC1 signaling modulates regulation of phosphate acquisition. An FDA-approved small-molecule inhibitor of Pho84 inhibits TORC1 signaling and potentiates the activity of the gold-standard antifungal amphotericin B and the echinocandin micafungin.

## Introduction

Organisms that fail to maximize growth in response to abundant nutrients can be outcompeted by those that do. Conversely, organisms that fail to cease growth and induce survival programs during stress and starvation lose viability. The Target of Rapamycin (TOR) signaling pathway is conserved in eukaryotes and integrates multiple channels of information regarding the cells’ nutritional and physical environment, to induce either growth and proliferation, or stress- and survival responses (1). In the human fungal pathogen *Candida albicans*, TOR participates in regulating morphogenesis (2–7) and responses to host cells (8). To control growth, *C. albicans* TOR also integrates signals of carbon source availability from the protein kinase A (PKA) pathway with nitrogen source status, its primary nutritional input (9). Similar TOR-PKA intersections have been reported in the model yeast *Saccharomyces cerevisiae* (10, 11).

In *S. cerevisiae*, TOR complex 1 (TORC1), which is susceptible to inhibition by rapamycin, is activated by preferred nitrogen sources such as glutamine and leucine. Leucine activates TORC1 through the Exit from G0 (EGO) complex by inducing GTP loading of one of its subunits, the small GTPase Gtr1 (12). Leucine also promotes TORC1 activity through leucine-tRNA synthase Cdc60, which physically interacts with Gtr1 (13). Many transcriptional regulators responsive to *S. cerevisiae* and *C. albicans* TORC1 pathways are conserved (4), but there are important differences between these species as well.

A small Ras-like GTPase upstream of TORC1, Rheb in mammals (14) and Rhb1 in *S. cerevisiae* (15), also responds to nutritional signals. Rheb is a central activator of mammalian TORC1 (mTORC1) and is modulated by the TSC1/TSC2 complex (16, 17), while in *S. cerevisiae*, Rhb1 seems to play a minor role in nutritional signaling. In *C. albicans*, Rhb1 is required for normal tolerance to rapamycin and for phosphorylation of ribosomal protein S6, a readout of TORC1 activation (9, 18). *C. albicans* Rhb1 co-regulates expression of genes important in nitrogen source uptake (18) and in virulence (19). Unlike *S. cerevisiae, C. albicans* also has a TSC2 homolog whose mutant phenotypes are consistent with a conserved GTPase-activating activity of *C. albicans* Tsc2 for Rhb1 (18).

Given the differences between the *S. cerevisiae* and *C. albicans* TORC1 pathways, we employed a forward genetic approach to find new components of *C. albicans* TORC1 signaling. Using our mariner transposon mutant collection (20), we isolated a rapamycin hypersensitive mutant in a *C. albicans* homolog of *PHO84*, the gene encoding the major *S. cerevisiae* high-affinity phosphate transporter.

Having identified a connection between *C. albicans* Pho84 and the *C. albicans* TOR pathway in a forward genetic screen, we characterized this link between phosphate homeostasis and the cell’s central growth control module. We found that we can indirectly target *C. albicans* TORC1, using small-molecule Pho84 inhibitors one of which is an FDA-approved antiviral drug, and that the antifungals amphotericin B and micafungin are potentiated by Pho84 inhibitors.

## Results

### A screen of haploinsufficient transposon mutants for altered rapamycin susceptibility identified a *PHO84* ortholog

We screened our heterozygous mutant collection of mariner-transposon insertions marked with our dominant selectable marker *NAT1* (20, 21) for altered rapamycin susceptibility. We isolated a transposon mutant hypersensitive to rapamycin in which the transposon disrupts the promoter of orf19.655, 67 bp upstream of the predicted translational start site (Fig. S1A). This orf encodes a protein with 66% amino acid identity to *S. cerevisiae* Pho84 and 55% amino acid homology to the *Piriformospora indica* PiPT phosphate transporter whose crystal structure was recently described (22) (Fig. S1B). According to Candida Genome Database (CGD) nomenclature, we called this orf *C. albicans PHO84*, and used the CGD sequence for further analysis (23). To confirm that rapamycin hypersensitivity of the transposon mutant was linked to the disrupted *PHO84* locus, two independent heterozygous deletion mutants and their homozygous null derivatives were constructed. Mutant phenotypes in these two lineages were the same, and one was chosen for further characterization (Fig. S1C). These mutants were also rapamycin hypersensitive (Fig. 1A), confirming that *PHO84* is required for normal tolerance of rapamycin.

**Fig. 1.**
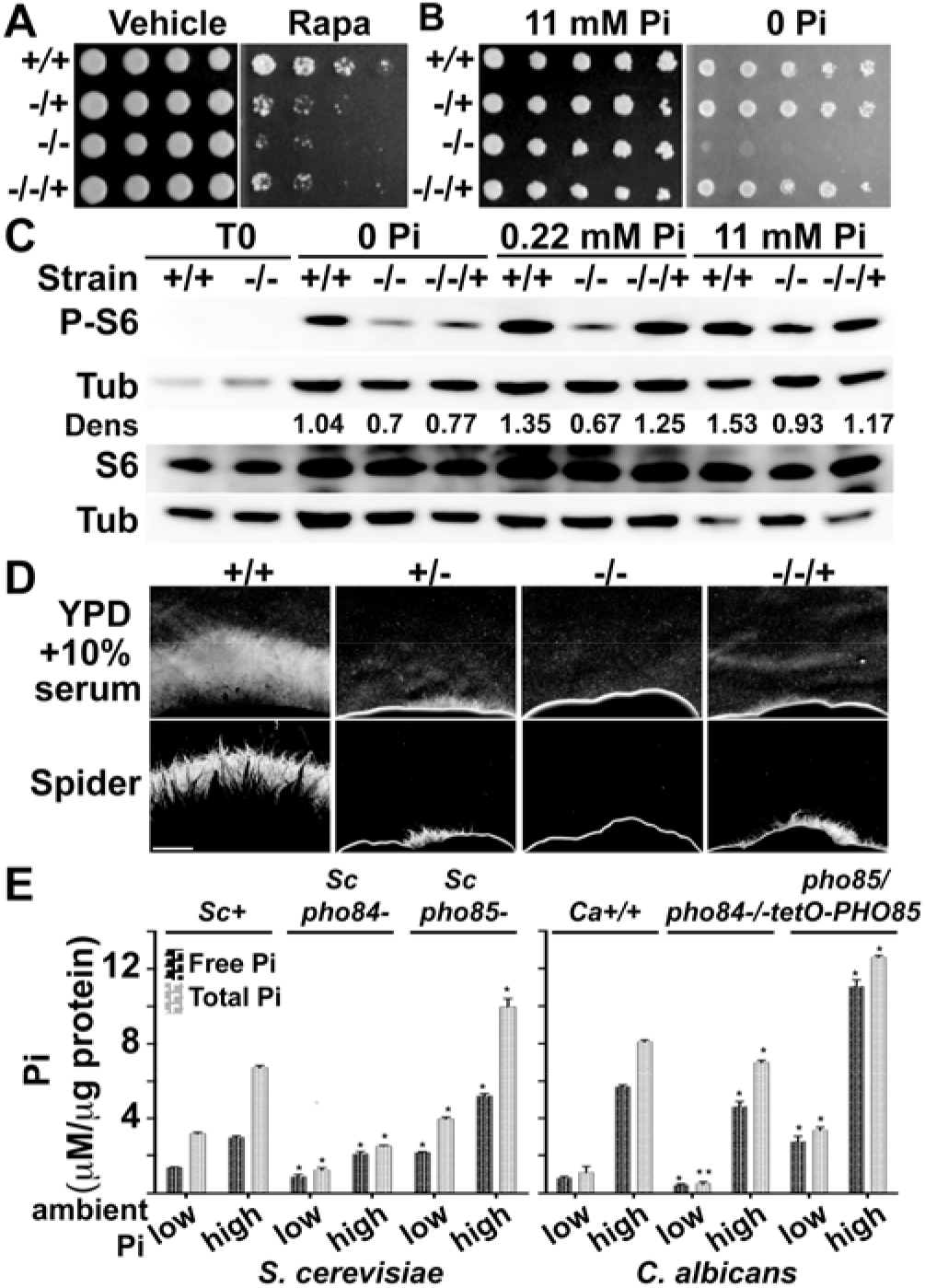
*C. albicans PHO84* is required for rapamycin tolerance, growth during Pi starvation, normal TORC1 activity and filamentation, and Pi homeostasis. **A**. Cell dilutions of wild type (wt) and a mutant series in *PHO84* pinned onto YPD with vehicle or 12 ng/ml rapamycin. *PHO84+/+*, JKC915. *pho84−/+*, JKC1583. *pho84−/−*, JKC1450. *pho84−/−/+*, JKC1588. **B**. Cells as in A pinned onto YNB with 0 or 11 mM Pi. **C**. Separate Western blots of the same samples for P-S6, total Rps6 and tubulin of wt (+/+, JKC915), *pho84* null (−/−, JKC1450), and *PHO84* reintegrant (−/−/+, JKC1588) cells grown in YNB with 0, 0.22 mM or 11 mM KH_2_PO_4_ for 90 min. Dens: densitometry of P-S6 vs. tubulin signal intensity. **D**. Strains as in (A) spotted at equidistant points around agar plates and spot edges imaged. Compare SI Fig. 2A. Bar 1mm. **E**. *S. cerevisiae* and *C. albicans* wild type and *pho84* null cells grown in SC medium with 0.22 mM (low Pi) or 11 mM Pi (high Pi) overnight, assayed for free and total Pi. *pho85* null cells as controls which hyperaccumulate Pi. *Sc+* (*S. cerevisiae PHO84 PHO85*), BY4741. *Scpho84−*, EY2960 and *Scpho85−*, from (37). Ca+/+ (*C. albicans PHO84/PHO84 PHO85/PHO85*), JKC915. *Capho84−/−*, JKC1450. *Capho85−/−*, CaLC1919, grown in 20 μg/ml doxycycline overnight. \**p*<0.01; \*\**p*<0.05. A-E represent at least 3 biological replicates; error bars SD of 3 technical replicates.

The cytoplasmic membrane protein Pho84 is the major high-affinity phosphate (Pi) transporter in *S. cerevisiae* (24–26). *PHO84* expression is controlled by the PHO regulon, a homeostatic system that maintains Pi availability for metabolism and growth in fluctuating external Pi conditions (27). *C. albicans pho84* mutants, like those in the *S. cerevisiae* homolog (28), failed to grow on medium without inorganic phosphate (Fig. 1B). Heterozygous and re-integrant cells, apparently haploinsufficient for rapamycin tolerance (Fig. 1A), grew robustly on this medium (Fig. 1B), indicating that mechanisms other than haploinsufficiency affect growth during Pi depletion, like the feedback loops between expression of high- and low-affinity Pi transporters characterized in detail in the *S. cerevisiae* PHO regulon (28). In liquid media with 1 mM Pi, growth of *pho84* cells was close to wild type (Fig S1D). Expression of the *C. albicans PHO84* homolog restored growth on medium lacking Pi to *S. cerevisiae pho84* mutants (Fig. S1E), indicating functional orthology. Wild type cells secrete acid phosphatase in response to low ambient Pi to mobilize covalently bound Pi from their environment, and this response was used for decades in studies of the *S. cerevisiae* PHO regulon (29). *C. albicans pho84−/−* mutants, like those in *S. cerevisiae* (24), inappropriately de-repressed acid phosphatase secretion in high ambient Pi (Fig. S1F), consistent with a conserved role of *C. albicans* Pho84 in the PHO regulon.

### *C. albicans* Pho84 is required for the normal TORC1 response to Pi availability

We then examined the relationship between Pho84 and TORC1. To test whether rapamycin hypersensitivity is due to decreased TORC1 kinase activity in the *pho84−/−* mutant, we monitored the phosphorylation state of ribosomal protein S6 (P-S6), which we previously showed is controlled by TORC1 signaling (9). Null mutants in *PHO84* had a weaker P-S6 signal than wild type during Pi refeeding at every Pi concentration of the media, though they responded to increasing Pi concentrations with an increasing P-S6 signal (Fig. 1C). Pho84 therefore is required for normal anabolic TORC1 signaling, and TORC1 activity responds to ambient Pi availability.

### Heterozygous and homozygous deletion mutants in *PHO84* are defective in hyphal morphogenesis

TORC1 regulates hyphal morphogenesis in *C. albicans* (2–7, 30), an important virulence determinant. Hyphal morphogenesis was defective in *pho84* mutants on YPD agar medium containing 10% serum, Spider medium and RPMI 1640 (Figs. 1D and S2A), while equal growth of mutant and wild type was seen in these media when the temperature signal for hyphal morphogenesis was absent (Fig. S2B). While many signaling pathways converge on morphogenesis, these findings are formally consistent with defective regulation by TORC1 (2).

### Pi content of cells lacking Pho84 is diminished

We asked if *C. albicans* TORC1 activity may be downregulated in response to decreased intracellular Pi in *pho84* mutants, analogously to the response of *S. cerevisiae* TORC1 to decreased intracellular amino acids. Using *pho85* mutants as controls known to hyperaccumulate intracellular Pi (31), we found that intracellular Pi concentrations were lower in *pho84−/−* null than in wild type cells in low and high Pi-containing media, though the difference was substantially less than in the homologous *S. cerevisiae* mutant-wild type pair (Fig. 1E). Diminished intracellular Pi concentrations of *C. albicans pho84−/−* cells may be responsible for, or contribute to the decreased TORC1 activation state, possibly in addition to lack of a putative TORC1-activating function performed specifically by Pho84.

### Gtr1 links Pho84 to TORC1 in *C. albicans*

Seeking a molecular link between Pho84 and TORC1 activity, we considered the possibility that Gtr1 may connect Pho84 to TORC1. *GTR1* was first described for its functional and physical proximity to *S. cerevisiae PHO84* (32, 33), and its product later was characterized as a component of the TORC1-activating EGO complex (12, 13, 34, 35). We found that the P-S6 response of *gtr1−/−* cells to phosphate refeeding was blunted (Fig. 2A). To determine whether this is an unspecific effect of decreased upstream TORC1 signaling, mutants in another small TORC1-activating GTPase, *RHB1*, were tested. The *rhb1* mutants responded to Pi refeeding like the wild type (Fig. 2B), suggesting that a Pi signal to TORC1 is transmitted specifically through Gtr1.

**Fig. 2.**
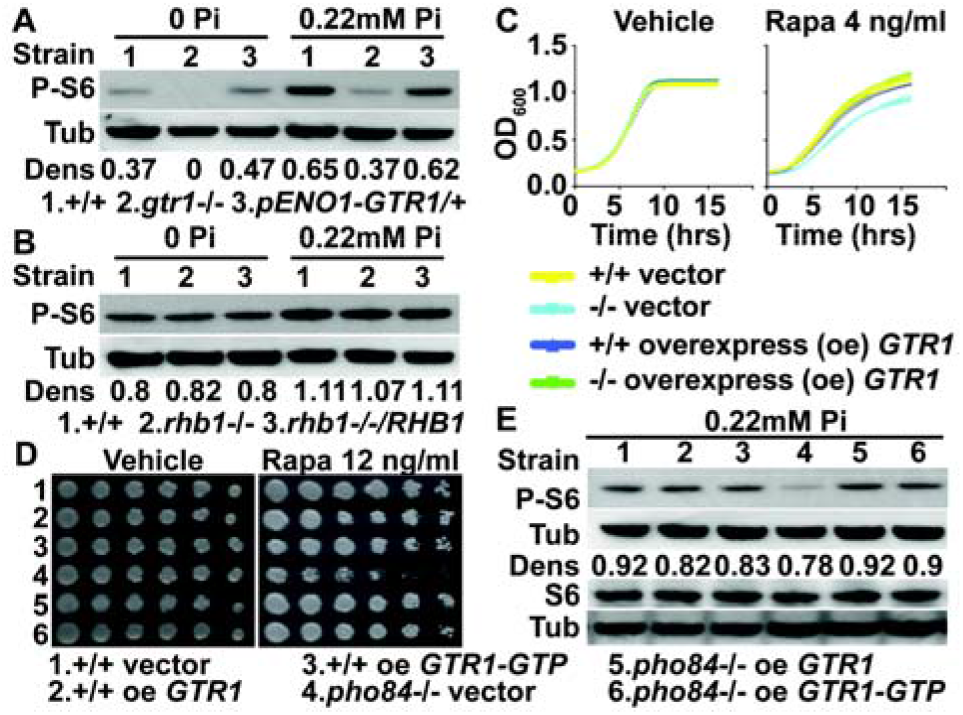
Gtr1 links Pho84 to TOR in *C. albicans*. **A**. Western blot of wild type (wt) (SC5314), *gtr1−/−* (YGM367) and *pENO1-GTR/GTR1* (YGM365) cells, grown in YNB with 0 and 0.22 mM KH2PO4 for 90 min. Dens: densitrometric ratio of P-S6 vs. tubulin signal intensity. **B**. Western blot of wt (SC5314), *rhb1−/−* (CCT-D1) and *rhb1−/−/pADH1-RHB1* (CCT-OE1) cells, grown in YNB with 0 or 0.22 mM KH_2_PO_4_ for 90 min. **C**. Growth in YPD with 4 ng/ml rapamycin or vehicle. OD_600_ monitored every 15 minutes. Yellow: wt with vector (JKC1594); blue: wt overexpressing *GTR1* (JKC1596); cyan: *pho84−/−* with vector (JKC1598); green: *pho84−/−* overexpressing *GTR1* (JKC1600). **D**. Cell dilutions pinned onto YPD with vehicle or 12 ng/ml rapamycin. Strains, (1) wt with vector (JKC1594), (2) wt overexpressing *GTR1* (JKC1596), (3) wt overexpressing GTR1-GTP (JKC1619), (4) *pho84−/−* with vector (JKC1598), (5) *pho84−/−* overexpressing *GTR1* (JKC1600) and (6) *pho84−/−* overexpressing GTR1-GTP (JKC1616). **E**. Western blot of cells grown in YNB with 0.22 mM KH_2_PO_4_ for 90 min. Strains1 -6 as in (D). A-E represent at least 3 biological replicates.

If Gtr1 acts downstream of Pho84 in activating TORC1, its overexpression may suppress *pho84−/−* phenotypes. *GTR1* was overexpressed from the *ACT1* promoter in wild type and *pho84−/− C. albicans* cells. Compared with rapamycin hypersensitive *pho84−/−* cells transformed with the empty vector, the resulting *pho84−/− pACT1-GTR1* cells showed wild type tolerance to rapamycin, suggesting recovery of their TORC1 signaling activity (Fig. 2C, D). To investigate this possibility TORC1 activity was tested directly by comparing the P-S6 signal of *pho84−/−* cells transformed with the empty vector with that of *pho84−/− pACT1-GTR1* cells. Overexpression of *GTR1* recovered Rps6 phosphorylation in *pho84−/−* cells nearly to wild type levels (Fig. 2E). A *GTR1* mutant encoding constitutively GTP-bound Gtr1^Q67L^, homologous to *S. cerevisiae* Gtr1^Q65L^ (12, 35, 36), was then constructed and overexpressed from the *ACT1* promoter. This GTR1-GTP allele suppressed the TORC1 signaling defect of *pho84−/−* cells apparently equally to the overexpressed wild type *GTR1* (Fig. 2D, E). Overexpression of *GTR1* or *GTR1-GTP* did not increase phosphorylation of S6 in wild type cells (Fig. 2E). These findings are consistent with the model that in *C. albicans*, Gtr1 indirectly or directly conveys a Pi signal to TORC1 and links Pho84 to TORC1 signaling.

We examined the relationship of Pho84 to TORC1 activity in the model yeast *S. cerevisiae*. A *pho84* null mutant in the S288C genetic background (37) was hypersensitive to rapamycin (Fig. S3A) at an intermediate ambient Pi concentration (1mM). Of note the rapamycin phenotype was highly responsive to the Pi concentration of pregrowth media. Rapamycin tolerance was not decreased by loss of *PHO84* in the Σ1278b background (not shown). Rapamycin hypersensitivity was not suppressed, but the *S. cerevisiae* Sch9 phosphorylation state (36) (Fig. S3B) and P-S6 signal intensity, which in *S. cerevisiae* also responds to TORC1 activation (Figs. 3 and S3C), were recovered by constitutively active Gtr1 in *pho84* null cells. These findings suggest that Pi homeostasis and TOR signaling are linked in *S. cerevisiae* as in *C. albicans*, though specific molecular connections seem to have divergently evolved in these two fungi.

**Fig. 3.**
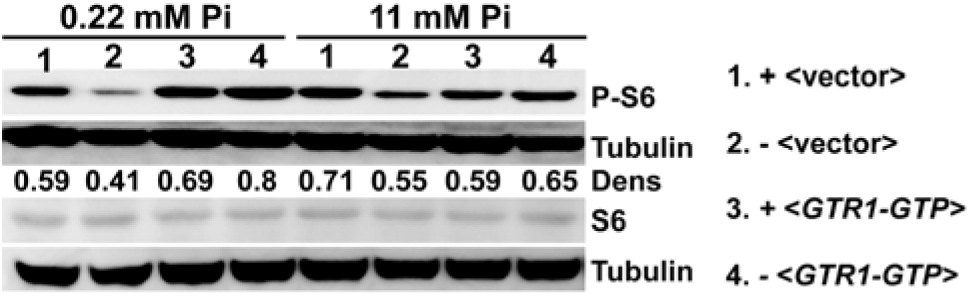
A connection between Pho84 and TORC1 is conserved in *S. cerevisiae*. Western blot of (1) *PHO84* with vector (Y1597), (2) *pho84* with vector (Y1599), (3) *PHO84* with *GTR1-GTP GTR2-GDP* (Y1596) and (4) *pho84* with GTR1-GTP *GTR2*-GDP (Y1604) cells, grown in SC (-His-Ura) containing 0.22 mM and 11 mM KH_2_PO_4_ for 90 min, probed for P-S6, S6 and tubulin. Representative of 3 biological replicates.

### TORC1 modulates the PHO regulon

As TORC1 not only responds to nutrient availability, but also directs nutrient uptake e.g. by regulating expression of amino acid and ammonium transporters, we questioned whether it may play a similar role in phosphate acquisition. Given known discrepancies between rapamycin exposure and physiological TOR modulation (38–41), we examined this potential connection genetically. Repressible *tetO* was used to control expression of *C. albicans TOR1* or of a hypomorphic *TOR1^Δ_1-381_^* encoding a protein lacking the first 381 amino acids which form protein-protein interaction HEAT repeat domains. The effect of *TOR1* depletion on expression of *PHO84* was then examined.

When wild type cells were transferred from overnight cultures into fresh rich medium, *PHO84* mRNA levels dropped, in accordance with the PHO regulon’s response to availability of fresh Pi sources. In cells depleted of either the wild type or the N-terminally truncated *TOR1* allele, *PHO84* expression also decreased but to a significantly lesser extent (Fig S4A). Full length *TOR1* permitted greater *PHO84* expression than the *TOR1^Δ_1-381_^* allele, suggesting structural perturbation of TORC1 by truncation of Tor1 affects its inhibitory as well as activating functions. Active TORC1, signaling nutritional repletion, hence contributes input to the PHO regulon to downregulate Pi starvation responses, while loss of TORC1 activity conveys a starvation signal to dampen these responses (Fig. S4A). Similarly, overexpression of Gtr1 and Gtr1-GTP blunted upregulation of secreted acid phosphatase in *pho84−/−* cells (Fig. S4B), supporting the model that in response to Pi TORC1 signaling downregulates the PHO regulon to integrate its activity with availability of other nutrients.

### Small-molecule inhibitors of Pho84 repress TORC1 and potentiate antifungal activity

*S. cerevisiae* Pho84 has been characterized as a Pi transceptor signaling to PKA, through identification of point mutations and small molecules that preferentially perturb transport, signaling or both (42, 43). Direct pharmacological inhibition of *C. albicans* TORC1 with rapamycin incurs too high a cost on host immune function to be clinically useful (44). We tested whether blocking Pho84 with its known small-molecule inhibitors phosphonoformic acid (foscarnet, Fos) and phosphonoacetic acid (PAA) (42), which we showed inhibit *C. albicans* growth in dependence on the presence of their target Pho84 (Fig. S5A, B), can indirectly inhibit *C. albicans* TORC1. Exposure of wild type cells to the FDA-approved antiviral foscarnet inhibited Rps6 phosphorylation (P-S6) in a dose-dependent manner (Fig. 4A), at Fos concentrations attained in human plasma during antiviral therapy (45, 46). In heterozygous cells (*pho84−/+*), whose haploisufficiency phenotypes likely reflect decreased copies of the drug target, Pho84, P-S6 was hypersensitive to Fos (Fig. 4A). In cells lacking the target Pho84, exposure to Fos did not further decrease the P-S6 signal (Fig. 4A). Pho84 inhibition with small molecules also recapitulated the hyphal growth defect seen in cells genetically depleted of *PHO84* (Figs. 1D, 4B and S5C).

**Fig. 4.**
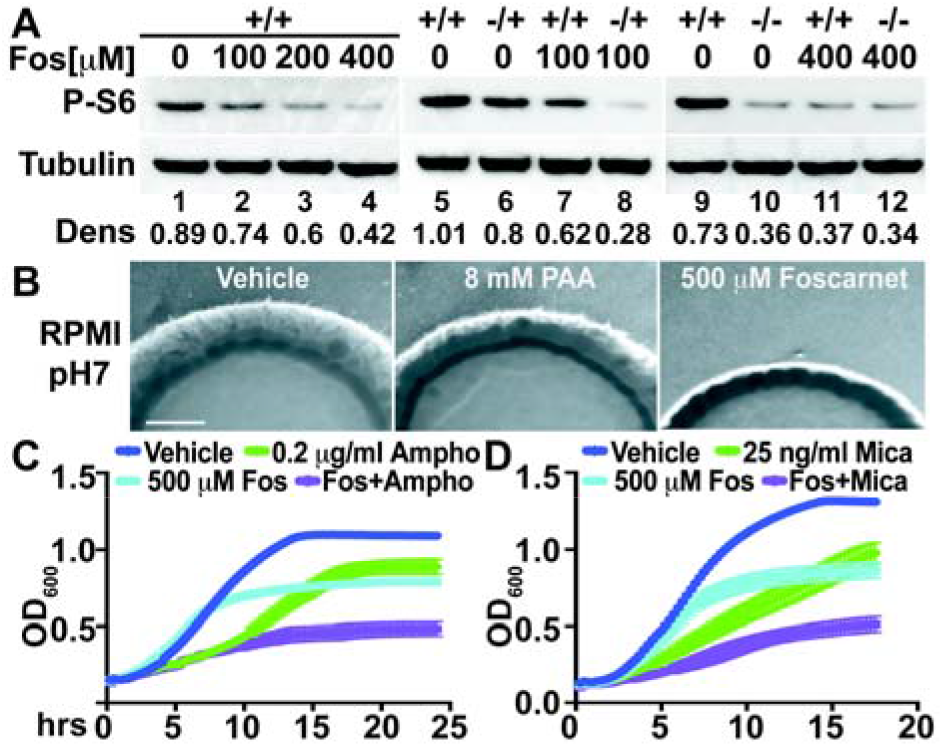
Small-molecule inhibition of Pho84 represses TORC1 and filamentous growth, and potentiates amphotericin B and micafungin. **A**. Western blot of wild type (wt) (SC5314) cells with (1) vehicle, (2) 100 μM foscarnet (Fos), (3) 200 μM Fos, (4) 400 μM Fos, and strains (5, 7, 9, 11) *PHO84+/+* (JKC915), (6, 8) *pho84−/+* (JKC1583) and (10, 12) *pho84−/−* (JKC1450), grown in standard SC (7.3 mM Pi) for 60 min, probed for P-S6 and tubulin. **B**. Wt (SC5314) spotted at equidistant points around RPMI agar (0.22 mM KH_2_PO_4_, pH7) containing vehicle, 8mM PAA, or 500 μM Fos, grown at 37°C, and spot edges imaged. Compare SI Fig. 2A. Bar 1mm. **C**. Wt (SC5314) exposed to vehicle, 500 μM Fos, 0.2 μg/ml amphotericin B, and amphotericin B plus Fos; OD_600_ in SC with 0.5 mM KH_2_PO_4_ at 30°C monitored every 15 minutes. **D**. Wt (SC5314) exposed to vehicle, 500 μM Fos, 25 ng/ml micafungin, and micafungin plus Fos; OD_600_ in SC with 0.5 mM KH_2_PO_4_ at 30°C monitored every 15 minutes.

Potentiating existing antifungals is a promising strategy (47–49). Fos at concentrations reached in plasma during antiviral therapy (45, 46), and PAA, potentiate activity of the antifungal amphotericin B, at concentrations of the latter far below those in serum or tissue during standard dosing (50) (Fig. 4C, S5D). Activity of the antifungal micafungin, belonging to the distinct drug class of echinocandins, was also potentiated (Fig. 4D). Because Pho84 is not conserved in mammals, inhibition of Pho84 offers a novel approach to fungal-specific TORC1 inhibition (Fig. 5) and to antifungal potentiation, as shown in our proof-of-principle experiments with PAA and Fos.

**Fig. 5.**
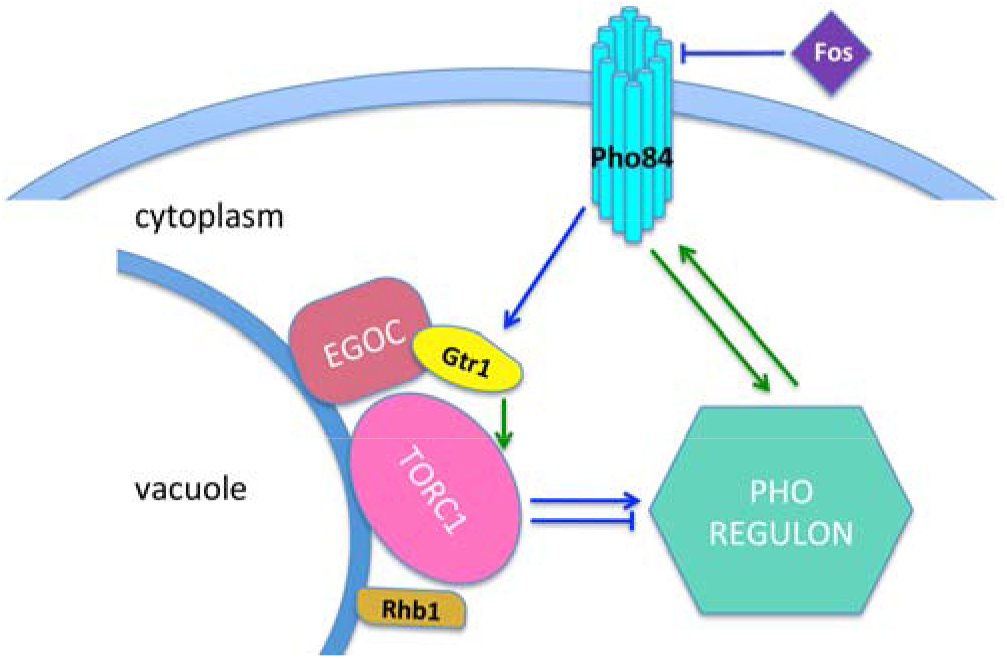
Pho84 activates TORC1 via Gtr1, and TORC1 in turn modulates the PHO regulon. Foscarnet inhibits Pho84 and thereby indirectly blocks TORC1 activity. Signaling events with known molecular mechanisms in *S. cerevisiae* are shown as green lines. Blue lines represent predicted activities based on our findings.

## Discussion

Eukaryotic cell mass consists to 0.5-1% of phosphorus (51). But phosphorus constitutes only 0.07% of the earth’s crust (51) and only ~15% of soil phosphorus is bioavailable in a soluble form (52). Fungi, like plants and bacteria, have sophisticated regulatory networks to manage Pi starvation. *S. cerevisiae* cells utilize the TORC1 and PKA pathways to ensure orderly cessation of growth when nitrogen or carbon sources become limiting (53, 54). *C. albicans* cells similarly use the TORC1 pathway for calibrated responses to distinct nitrogen sources, with modulating input from PKA according to carbon source type and concentration (9).

Our study shows that *C. albicans* TORC1 monitors not only nitrogen and carbon source-but also Pi availability (Fig.1C; 2A,E). Whether Pho84 activates TORC1 indirectly by an increased intracellular Pi content, directly in a transceptor role, or through a combination of these inputs, remains an open question. Our results establish the small GTPase Gtr1, first discovered in *S. cerevisiae* due to its mutant phenotypes’ resemblance to those of Pho84 (32), as an important element linking *C. albicans* Pho84 and TORC1 (Fig. 2). How Gtr1, a subunit of the TORC1-associated vacuolar membrane-residing EGO complex (55), receives information from Pho84 remains to be determined. Possible models include an activating signal to the TORC1-activating SEACAT complex (55) through a sensor of cytosolic Pi or vacuolar polyphosphate, or a physical interaction between Pho84 and the EGO complex on endosomes while Pho84 is internalized during ambient Pi abundance (56). We cannot exclude alternative explanations of our findings, e.g. a TORC1 response only to intracellular Pi concentrations, and upregulation of low-affinity Pi transporters during *GTR1* overexpression in *pho84−/−* null cells. If Gtr1 is activated by Pho84, this GTPase’s role in the input to TORC1 regarding the cell’s Pi state appears to be specific (Fig. 2), as there was no perturbation of TORC1 activation during Pi refeeding in *rhb1−/−* cells (Fig. 2B).

In *S. cerevisiae*, Pho84 signals to PKA as a transceptor (42, 57). A signal from Pho84 to TORC1 has not yet been described in *S. cerevisiae*. We found that in the genetic background S288C loss of Pho84 leads to rapamycin hypersensitivity and TORC1 inactivation as it does in *C. albicans* (Fig. S3). Sch9 phosphorylation, like that of Rps6, is known to correspond with the TORC1 activation state (36, 58). Overexpression of constitutively active Gtr1 restores TORC1 activity in *S. cerevisiae*, as assayed by Sch9 and Rps6 phosphorylation (Fig. 3, S3B). But in contrast to *C. albicans*, constitutively active Gtr1 did not suppress rapamycin hypersensitivity of *S. cerevisiae pho84* null cells, indicating differences in the connections of PHO and TORC1 signaling between these fungi. In *S. cerevisiae*, control of entry into quiescence (G0) by the Rim15 kinase is coregulated by the PHO pathway cyclin/cyclin-dependent kinase module Pho80/Pho85, as well as by TORC1(59), suggesting multiple levels of cross-talk between phosphate-specific and global nutritional signaling pathways in that organism, which remain to be explored in *C. albicans*.

In addition to the signal from Pho84 to TORC1, we observed a signal in the opposite direction, from TORC1 to expression of *PHO84* and to the classic readout of the PHO regulon, the acid phosphatase (27) (Fig. S4A), though clearly input from TORC1 contributes to only a fraction of the PHO regulon responses. TORC1 is well known to regulate proteins required for acquisition of other nutrients like amino acids and ammonium in *S. cerevisiae* (60, 61). TORC1 input to the PHO regulon may fine-tune the investment of energy and nutrients on Pi acquisition to match the overall state of the cell. In *S. cerevisiae*, transcriptional regulation favoring Pi acquisition is achieved through the transcription factor Pho4 (31). A *C. albicans* homolog of Pho4, required for stress resistance (62, 63) and commensalism in a murine model (63), was recently shown to control *C. albicans PHO84* expression (62). How *C. albicans* Pho4 may be co-regulated by TORC1 remains to be determined.

While TORC1 monitors nutrient availability in mammals as in fungi, its relationship with the PHO regulon may have diverged in these phyla. In fungi, H^+^-Pi symport is the major form of Pi import, since the steep Pi concentration gradient at the plasma membrane imposes a high energetic demand on transport, met by the electrochemical proton gradient generated by the P-type H^+^ ATPase (52). In animals, Na^+^-Pi pumps predominate (64). Humans largely consume Pi together with amino acids in food consisting of other eukaryotes and their products, so that Pi starvation tends to occur during starvation for protein (65). In contrast, Pi and nitrogen sources must be acquired independently by unicellular organisms, so the connection we discovered between the fungal H^+^-Pi symporter and the TORC1 pathway appears physiologically plausible.

We found that TORC1 is indirectly repressed during Pho84 inhibition with small molecules (Fig. 4, S5). Since TORC1 components are conserved between fungi and humans while Pho84 has no human homologs, this result provides an option for indirect, fungal-specific TORC1 inhibition not previously explored in the search for new antifungals. While TORC1 is only partially inhibited in this manner, other fungal-specific indirect TORC1 activators may provide synergistic targets. More immediately, Pho84 inhibitors potentiate the antifungal effect of micafungin and of amphotericin B (Fig. 4C,D), possibly permitting lower dosing of the latter, “gold-standard” broad-spectrum agent to obviate its often treatment-limiting toxicities (66). We used 2 of multiple Pho84 inhibitors previously characterized (42) (Fig. 4 and S5D), and more specific, non-competitive inhibitors with more favorable therapeutic indices than Fos may be found through screening efforts. Since Pho84 homologs are highly conserved among fungi, potentiating amphotericin B activity through their inhibition may prove a viable therapeutic strategy for other fungal species less amenable to other antifungal agents.

## Materials and Methods

### Strains and culture conditions

The *C. albicans* and *S. cerevisiae* strains, plasmids and primers used are described in SI Tables 1-3. *C. albicans* strains were generated in the SN genetic background using *HIS1* and *ARG4* markers (67), as well as the *CaNAT1* selectable marker, as described in (20, 21). Two independent heterozygotes were used to derive homozygous null and reintegrant mutants of *PHO84*, as well as *tetO-TOR1* mutants. Auxotrophies were complemented so that only prototrophic strains, or strains with identical auxotrophies, were compared in an experiment. Introduced mutations were confirmed by PCR spanning the upstream and downstream homologous recombination junctions of transforming constructs, and sequencing. Experiments with defined ambient Pi concentrations were performed in YNB 0 Pi (ForMedium Ltd, Norfolk, UK) with added KH_2_PO_4_ to stated concentrations. Other media were used as in (20).

### Screening transposon mutants for altered rapamycin susceptibility

Our heterozygous mutant collection containing a mariner transposon marked with *CaNAT1* (20) was used. Mutants were replicated to a 96-well plate containing 2xYPD with 8% glucose and grown to saturation at room temperature, to minimize filamentous growth. Cells were replicated to YPD agar medium with vehicle (90% ethanol) or 20 ng/ml rapamycin. Clones showing less growth than the wild type (SC5314) were isolated as rapamycin hypersensitive. The transposon insertion site was identified by vectorette PCR as previously described (20).

### Growth Assays

For cell dilutions spotted onto agar media as previously described (20), saturated overnight cultures were diluted in 5-fold steps from an OD_600_ of 0.5. For growth curves in liquid media, saturated overnight cultures in YPD were washed once in 0.9% NaCl and diluted to an OD_600_ of 0.15 in 150 μl medium in flat bottom 96-well dishes. For growth assays including those during drug exposure, OD_600_ readings were obtained every 15 min in a plate reader and standard deviations were calculated and graphed in Graphpad Prism. Growth during drug exposure was assayed in SC medium. Vehicle for Pho84 inhibitors PAA (Sigma, 284270) and Fos (Santa Cruz Biotechnology, SC-253593A) was water and for amphotericin B (Sigma, A9528), DMSO.

### Western Blots

Cell harvesting, lysis and Western blotting were performed as described in (9). Antibodies are listed in SI Table 1. At least three biological replicates were obtained. For densitometry, ImageJ (imagej.net/welcome) software (opensource) was used to quantitate signals obtained on a KODAK Image Station 4000MM.

### Filamentous growth assay

Cells were revived from frozen stocks on solid YPD overnight, washed and resuspended in 0.9% NaCl to OD_600_ 0.1. Variations between single colonies and colony density effects were minimized by spotting 3 μl cell suspension at 4 or 6 equidistant points, using a template, around the perimeter of an agar medium plate, which then were incubated and imaged as in (20). For small-molecule Pho84-inhibitor effects on filamentation, Spider and RPMI was used (TOKU-E, Cat# R8999-04), the latter with 0.22mM KH_2_PO_4_ buffered to pH 7 with 50mM MOPS. At least 3 biological replicates were obtained for each condition.

### Acid phosphatase assays

As adapted from (25), overnight cultures in SC were diluted to an OD_600_ 0.05 into YNB medium buffered to pH4 with 50mM sodium citrate containing 0 or 11mM KH_2_PO_4_ and grown overnight. They were washed thrice with water, and p-Nitrophenyl Phosphate (Sigma, N4645) was added to a concentration of 5.62mg/ml. After 15 mins at room temperature the reaction was stopped with Na_2_CO_3_ (pH=11) to a concentration of 0.3g/ml, and OD_420_ and OD_600_ were measured. At least 3 biological replicates, with 3 technical replicates each, were obtained.

### Intracellular Pi assays

Free and total Pi was measured by colorimetric molybdate assay as described (68). Briefly, cultures were washed with distilled water twice, resuspended in 500 μL 0.1% Triton X-100, and lysed by glass bead homogenization. Lysate protein content was determined using a BioRad Protein Assay kit. Free Pi was measured in unboiled lysate, then total phosphate was measured after boiling 3–30 μg of whole cell lysate for 10 min in 1 N H_2_SO_4_. At least 3 biological replicates, with 3 technical replicates each, were obtained.

### RT-PCR expression analysis

Cells were grown overnight in YPD medium with 5ng/ml doxycycline, diluted into YPD with 30 μg/ml doxycycline, and harvested at time 0, 2 h and 4 h. RNA was extracted with the Direct-Zol RNA miniprep kit (Zymo Research # R2051). RT-PCR procedures were performed as indicated (69).

## Acknowledgments

We thank Gerald R. Fink, Erin K. O’Shea, Bin He, Charles Boone, Françoise Stutz, Kevin Struhl and Fred Winston for plasmids and *S. cerevisiae* strains; Leah Cowen, Chung-Yu Lan and Rajini Rao for *C. albicans* strains; Gerald R. Fink, Felix Lam, Valmik Vyas, Luke Whitesell and Bin He for helpful conversations; Brian M. Hoffman, Valeria C. Culotta and Amit R. Reddi for protocols; and Dennis Wykoff, Bin He, Robert Husson and Paula Watnick for critical reading of the manuscript. This work was funded by NIAID R21AI096054 and R01AI095305. P.F. was supported by Science Foundation Ireland Grant 11/RFP.1/GEN/3044. M.E.C. was funded by NCI R01 CA154499.

